# GABA administration prevents severe illness and death following coronavirus infection in mice

**DOI:** 10.1101/2020.10.04.325423

**Authors:** Jide Tian, Blake Middleton, Daniel L. Kaufman

**Author notes:** Corresponding author: Daniel Kaufman, Department of Molecular and Medical Pharmacology, David Geffen School of Medicine at the University of California, Los Angeles, CA. 90095-1735. or.

## Abstract

There is an urgent need for new treatments to prevent and ameliorate severe illness and death induced by SARS-CoV-2 infection in COVID-19 patients. The coronavirus mouse hepatitis virus (MHV)-1 causes pneumonitis in mice which shares many pathological characteristics with human SARS-CoV infection. Previous studies have shown that the amino acid gamma-aminobutyric acid (GABA) has anti-inflammatory effects. We tested whether oral treatment with GABA could modulate the MHV-1 induced pneumonitis in susceptible A/J mice. As expected, MHV-1-inoculated control mice became severely ill (as measured by weight loss, clinical score, and the ratio of lung weight to body weight) and >60% of them succumbed to the infection. In contrast, mice that received GABA immediately after MHV-1 inoculation became only mildly ill and all of them recovered. When GABA treatment was initiated after the appearance of illness (3 days post-MHV-1 infection), we again observed that GABA treatment significantly reduced the severity of illness and greatly increased the frequency of recovery. Therefore, the engagement of GABA receptors (GABA-Rs) prevented the MHV-1 infection-induced severe pneumonitis and death in mice. Given that GABA-R agonists, like GABA and homotaurine, are safe for human consumption, stable, inexpensive, and available worldwide, they are promising candidates to help prevent severe illness stemming from SARS-CoV-2 infection and other coronavirus strains.

---

While GABA is well known as a neurotransmitter that is commonly used in the central nervous system, it is becoming increasingly appreciated that many immune cells express GABA-Rs. The biological roles of GABA-Rs on immune cells are not yet well understood, but there is a growing body of evidence that the activation of these receptors generally has immunoregulatory actions. In the innate immune system, antigen-presenting cells (APCs) express GABA-A-type receptors (GABA_A_-Rs) which form a chloride channel and their activation reduces APC reactivity (1, 2). Neutrophils express GABA-B-type receptors (GABA_B_-Rs) which are G-protein coupled receptors that modulate their function (3). In the CNS, microglia express both GABA_A_-Rs and GABA_B_-Rs and their activation reduces microglia responsiveness to inflammatory stimuli (4). Alveolar macrophages express GABA_A_-Rs and application of a GABA_A_-R-specific agonist decreases the expression of many pro-inflammatory molecules in cultures of LPS-stimulated lung macrophages (5). In the adaptive immune system, we have shown that GABA-R activation promotes effector T cell cycle arrest without inducing apoptosis (6). *In vivo*, administration of GABA, or the GABA_A_-R-specific agonist homotaurine, inhibits autoreactive Th1 and Th17 cells while promoting CD4^+^ and CD8^+^ Treg responses (7-9). Taking advantage of these properties, we have demonstrated that administration of GABA or homotaurine inhibits disease progression in mouse models of type 1 diabetes (T1D), multiple sclerosis, and rheumatoid arthritis, and limits inflammation in a mouse model of type 2 diabetes (1, 6, 8-10).

There is an urgent need to develop new treatments to reduce severe illness and death in COVID-19 patients. Patients who develop severe illness appear to mount weaker and delayed innate immune responses to SARS-CoV-2 infection, which leads to excessive adaptive immune responses later that do not taper off appropriately (11-13). This can lead to “cytokine storms”, disseminated intravascular coagulation, multiple organ dysfunction syndrome (MODS), and death. Studies of anti-CD3-activated human PBMC have shown that GABA inhibits IL-6, CXCL10/IP-10, CCL4, CCL20, and MCP-3 production (14). Longitudinal studies of COVID-19 patients reveal that high levels of serum IL-6 and Th1, Th17, and Th2-secreted proteins are associated with progression to severe illness (11, 15). Many of these biomarkers of severe illness have been shown to be reduced by GABA-R agonists in the aforementioned *in vitro* studies of human PBMC and/or mouse models of autoimmune disease. Currently, however, there is no information on whether GABA treatment modulates the outcome of viral infections.

Like SARS-CoV and SARS-CoV-2, mouse hepatitis virus (MHV)-1 is a pneumotropic beta-coronavirus of the group 2 lineage and is widely used as a safe model of SARS-CoV infection (16-19). MHV-1 infection creates a lethal pneumonitis, similar to SARS-CoV-induced disease, in A/J mice. Intranasal inoculation with 5000 plaque-forming units (PFU) of MHV-1 in A/J mice induces an acute respiratory distress syndrome with a high lethality rate. The infected mice develop pathological features of SARS-CoV-2, including high levels of pulmonary cytokines/chemokines, pneumonitis, dense macrophage infiltrates, hyaline membranes, fibrin deposits, accompanied by loss of body weights and respiratory distress (16-19).

We studied whether oral GABA treatment beginning at the time of MHV-1 inoculation or starting three days post-inoculation (by which time signs of illness are apparent), could modulate the severity of the ensuing illness and the rate of death

## Materials and methods

### Mice

Female A/J mice (7 weeks in age) were purchased from the Jackson Laboratory and maintained in microisolator cages and fed with a standard diet and water *ad libitum*. One week after arrival, they were inoculated with MHV-1. The mice were immediately randomized and treated (or not treated) with GABA, as described below. This study was carried out in accordance with the recommendations of the Guide for the Care and Use of Laboratory Animals of the National Institutes of Health. The protocols for all experiments using vertebrate animals were approved by the Animal Research Committee at UCLA.

### Reagents

GABA was purchased from Millipore-Sigma (stock # A2129, St. Louis, MO, USA).

### Virus

MHV-1, DBT cells, and HeLa-CECAM1 were generously provided by Dr. Stanley Perlman (University of Iowa). MHV-I virus was prepared and titered as previously described (16-19).

### Viral infection and GABA treatment

At 8 weeks in age, female A/J mice were anesthetized and inoculated intranasally with 5000 PFU MHV-1 in 50 μl cold Dulbecco’s modified Eagle’s medium (DMEM). The mice were immediately randomized and provided with plain water (controls) or water that contained GABA (20 mg/ml) for the entirety of the observation period. Another group of MHV-1 inoculated mice received plain water for three days, by which time they displayed signs of illness, and then were placed on GABA-containing water for the rest of the observation period. We monitored their body weights daily beginning on the day of infection and up to 14 days post-infection.

### Illness scoring

Individual mice were monitored for illness development and progression which were scored on the following scale: 0) no symptoms, 1) slightly ruffled fur and altered hind limb posture; 2 ruffled fur and mildly labored breathing; 3) ruffled fur, inactive, moderately labored breathing; 4) ruffled fur, obviously labored breathing and lethargy; 5) moribund and death.

The percent survival of each group of mice was determined longitudinally for each group. Mice with a disease score of 5 were weighed, euthanized, and their lungs removed and weighed for calculation of lung coefficient index (the ratio of lung weight to total body weight, which reflects the extent of edema and inflammation in the lungs). On day 14 post-infection, the surviving animals were weighed, euthanized, and their lungs were removed and weighed for determination of the lung coefficient index.

## Results and Discussion

Following MHV-1 inoculation, the mice receiving plain water began to progressively lose body weight each day. By day 6, this control group had lost an average of 23% of their weights, as expected (16-19). At this time point, the mice that had been given GABA immediately after MHV-1 infection had lost an average of 11% of their body weights, and those given GABA three days after infection had lost an average of 17% of their body weights (Fig. 1A). After day 6, the mice in the control group began to succumb to their illness and only 3/9 mice survived to day 14 post-infection (Fig. 1B). In contrast, none of the mice given GABA starting immediately after MHV-1 inoculation died (Fig. 1B), and their body weight was on average only 7% below their starting weights 14 days post-infection (Fig. 1A). Of the mice that began GABA treatment 3 days post-infection, 1/9 mice died (on day 9), and their body weights at 14 days post-infection were 90% of their starting weight. The survival curves for each group are shown in Fig. 1B.

**Figure 1.**
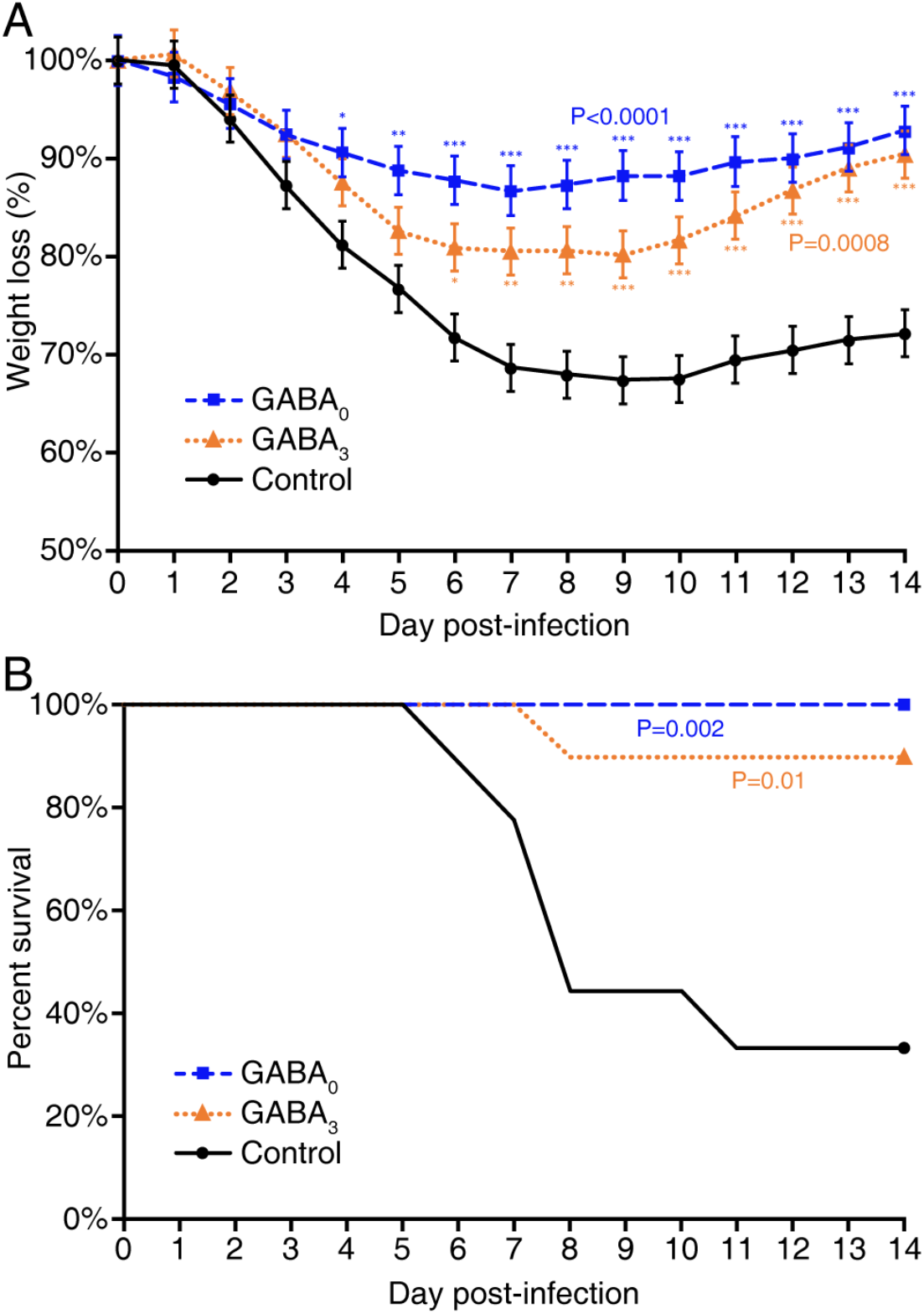
GABA treatment reduces body weight loss and death rate in MHV-1 infected mice. Female A/J mice were inoculated with MHV-1 intranasally and immediately placed on plain water (control, solid black line) or water containing 20 mg/ml GABA (GABA_0_, dashed blue line) or given plain water for 3 days post-infection and then placed on water containing 20 mg/ml GABA (GABA_3_, orange dotted line) for the remaining observation period. **A)** Daily changes in % body weights post-infection (% of day 0), p<0.0001 and p=0.0008 for GABA_0_ and GABA_3_ (respectively) vs. control by repeated measure ANOVA. (GABA_0_ vs. GABA_3_,p=0.175.) **B)** Daily percent of surviving mice in each group, p=0.002 and p=0.01 for GABA_0_ and GABA_3_ vs. control, respectively by log-rank test. (p=0.32 for GABA_0_ vs. GABA_3_) N=9 mice in the control group, 10 mice in each GABA-treated group. Data shown are from two separate studies with 4-5 mice/group. *p<0.05, **p<0.01, ***p<0.001 vs. the control.

**Fig. 2.**
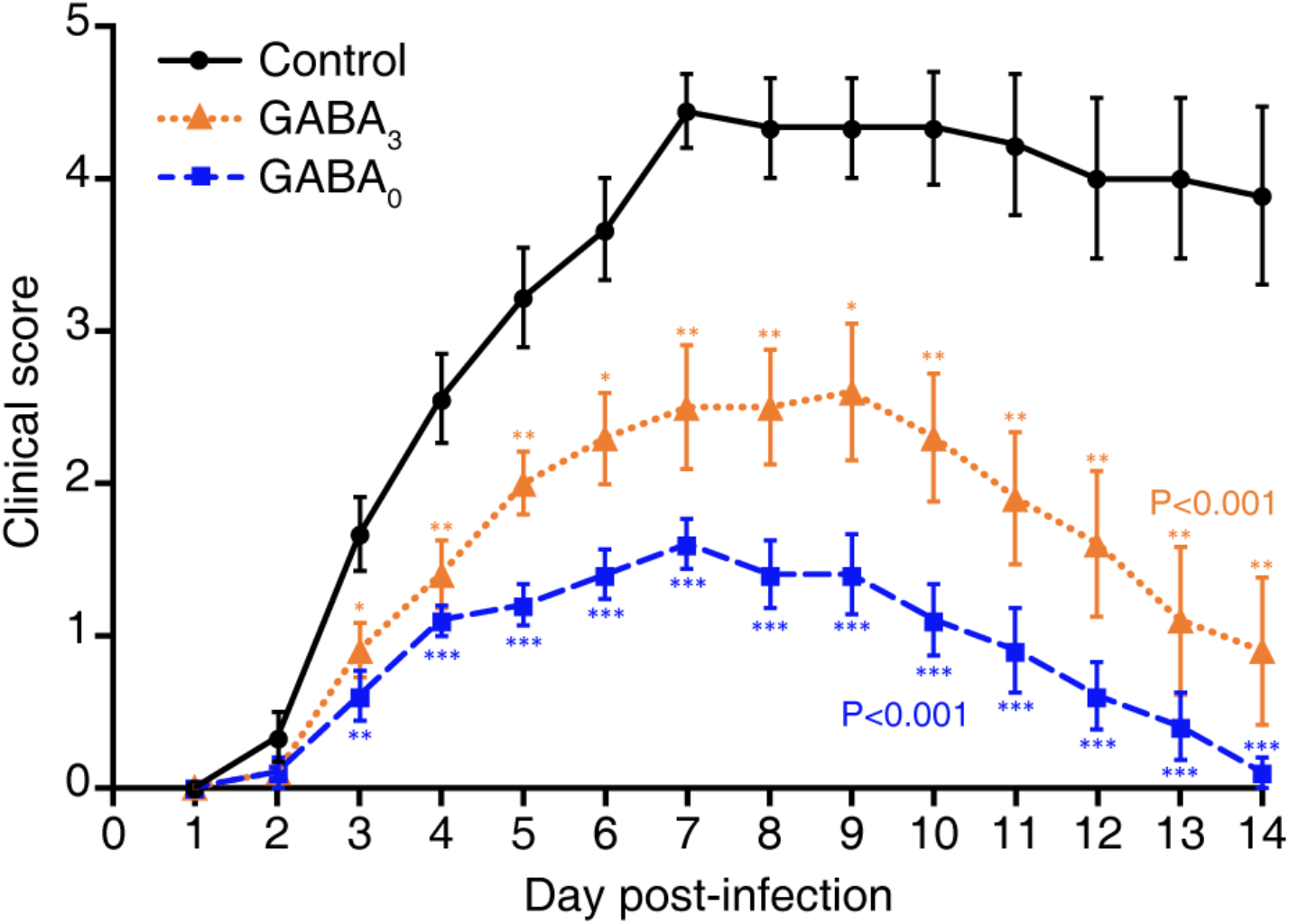
GABA treatment reduces illness scores in MHV-1 infected mice. The animals described in Fig. 1 were scored daily for the severity of their illness as detailed in Methods. Data shown are the mean clinical scores +/-SEM of each group from two separate experiments. Overall p<0.001 for GABA_0_ and GABA_3_ vs. control. GABA_0_ vs. GABA_3_ p=0.042 using the Kruskal-Wallis test. *p<0.05, **p<0.01, ***p<0.001.

**Figure 3.**
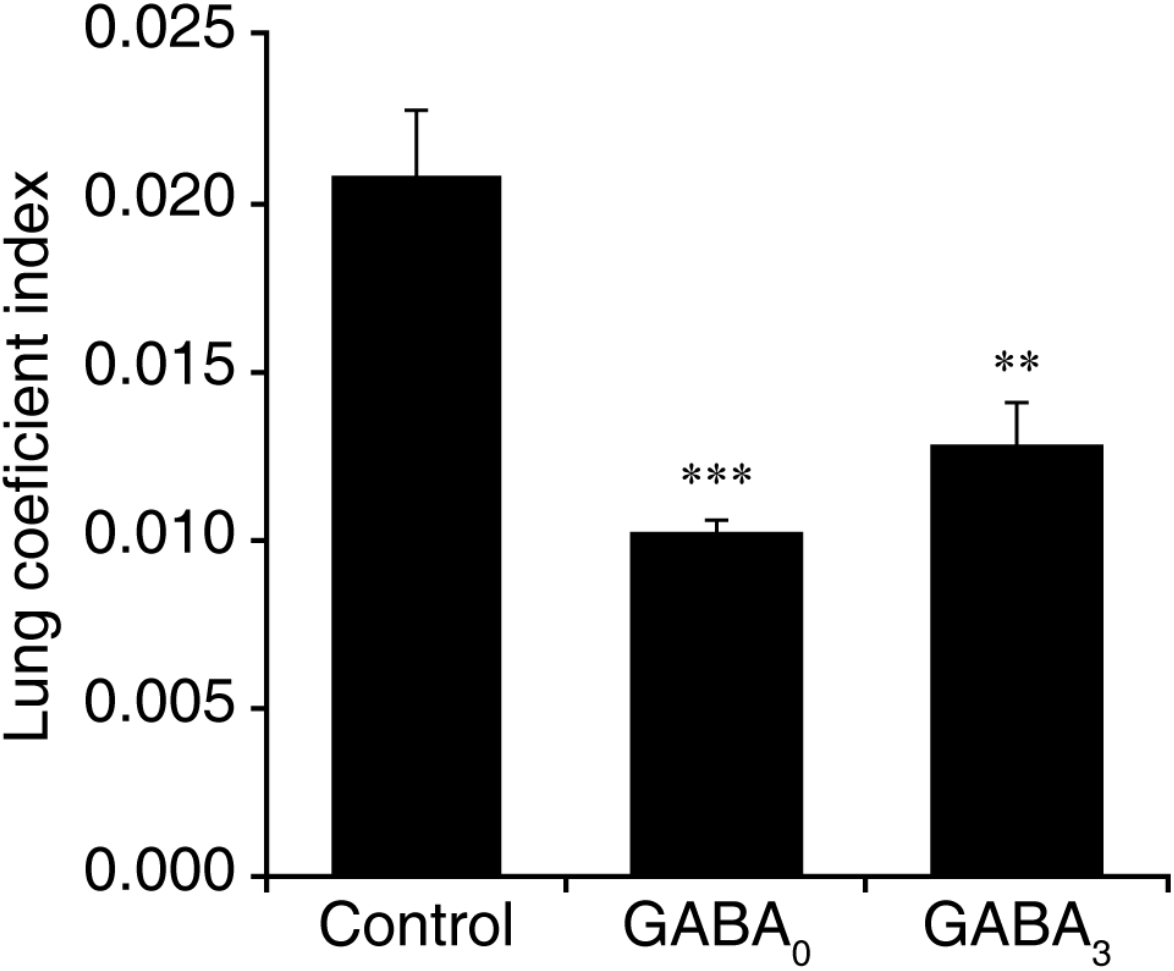
GABA treatment reduces the lung coefficient index in MHV-1 infected mice. The lungs were harvested and weighed when an animal became moribund or at 14 days post-infection. Data shown are the mean lung coefficient index ± SEM for each group from two separate studies. ***p<0.001 and **p<0.01 for GABA_0_ and GABA_3_ (respectively) vs. control water treated group by Student’s t-test.

In terms of illness, MHV-1 infected control mice began to display signs of illness two days post-infection and rapidly became severely ill thereafter, with their illness peaking around day 7 post-infection. While most control mice died between days 6-11 post-infection, those that survived displayed only partial recovery from illness. In contrast, the mice receiving GABA immediately after MHV-1 inoculation developed only mild illness, with the highest average illness score of 1.6 on day 7 post-infection. Illness in the mice given GABA at 3 days post-infection was also significantly reduced compared to that in the control group, and their maximum mean illness score was 2.5. Thus, GABA treatment immediately after MHV-1 infection, or 3 days later when the clinical signs of the disease were apparent, reduced the subsequent severity of the disease and the death rate.

The lung coefficient index reflects the edema and inflammation in the lung. The lung coefficient index of mice that were given GABA immediately after MHV-1 infection was 49% of that of control mice (p<0.001). The mice receiving GABA treatment beginning 3 days post-infection had a lung coefficient index that was 62% of that in the control mice (p<0.01). This provides an independent measure indicating that GABA treatment limited the MHV-1 induced pulmonary edema and inflammation in A/J mice.

Together, the reduction in body weight loss, illness scores, death rate, and lung coefficient index indicate that GABA treatment can reduce illness severity and death rate following coronavirus infection, even when the treatment is initiated after symptoms appear.

Since weaker and delayed early immune responses to SARS-CoV-2 infection are associated with more severe illness in COVID-19 patients (11) and GABA has anti-inflammatory effects, we anticipated that treatment with GABA immediately after MHV-1 infection might be deleterious by limiting or delaying innate immune responses. We were surprised that early GABA treatment immediately after MHV-1 infection was very effective in preventing illness progression and death, suggesting a rapid effect of GABA on innate immune responses and/or the lung airway cells. The lung epithelial cells of mice and humans also express GABA_A_-Rs (20, 21). It is possible that the activation of these GABA_A_-Rs leads to Cl^-^ efflux, which would act to limit Ca^2+^ influx in these epithelial cells. Because many viruses, including coronaviruses, elevate intracellular Ca^2+^ concentrations in order to enhance viral replication (22, 23), the activation of GABA_A_-Rs may have limited MHV-1 replication, a possibility that we are currently investigating.

Additionally, treatment with GABA, a GABA_B_-agonist, or GABA_A_-R positive allosteric modulators can reduce inflammation and improve alveolar fluid clearance and lung functional recovery in rodent models of acute lung injury (24-28). Whether these capabilities contributed to GABA’s ability to limit the progression of pneumonitis soon after MHV-1 infection is an open question. Thus, treatment with GABA-R agonists may have multiple beneficial actions for the treatment of COVID-19 patients.

GABA treatment was tested in hundreds of epilepsy patients for its ability to reduce seizures (29-31). While it had no clinical benefit (probably because it cannot cross the blood-brain barrier), it had no adverse effects in these long-term studies. A more recent phase Ib GABA oral dosing study also indicated that GABA is safe (32) and there are currently several ongoing clinical trials that are administering oral GABA to individuals with T1D (ClinicalTrials.gov Identifiers: NCT02002130, NCT03635437, NCT03721991, NCT04375020). In addition, the GABA_A_-R specific agonist homotaurine was tested in a large long-term phase III clinical trial for Alzheimer’s disease, and while it was not effective it had an excellent safety record (see (8, 9) for a discussion of homotaurine’s safety). Both GABA and homotaurine are inexpensive, stable at room temperature, and available worldwide making them excellent candidates for clinical testing as adjunctive treatments for COVID-19.

These studies indicate the potential usefulness of GABA and/or homotaurine as a treatment for COVID-19 and other coronavirus infections. However, until clinical trials are completed and GABA and/or homotaurine are approved for use in the treatment of COVID-19 by relevant governing bodies, GABA and homotaurine should not be consumed by COVID-19 patients as they may pose health risks, such as dampening beneficial immune or physiological responses.

## Acknowledgments

We would like to thank Dr. Stanley Perlman for generously providing MHV-1, DBT cells, and HeLa-CECAM1 cells. We would also like Drs. Min Song for assistance, Cindy Chau for advice, and Jeffery Gornbein for statistical analysis. This work was supported by grants to DLK from the UCLA DGSOM-Broad Stem Cell Research Center COVID-19 Research Award (ORC #20-34), The National Institutes of Health (R21 DE029020), and the Department of Defense (CDMRP PR191176), as well as DLK’s unrestricted funds.

## Author Contributions

Conceived and designed the experiments: JT, DLK; Performed the experiments: JT, BM; Analyzed the data: JT, DLK. Wrote the paper: JT, DLK. Drs. Daniel Kaufman and Jide Tian are guarantors of this work and, as such, had full access to all the data in the study and take responsibility for the integrity of the data and the accuracy of the data analysis.

